# Epigenomic landscape of the human dorsal root ganglion: sex differences and transcriptional regulation of nociceptive genes

**DOI:** 10.1101/2024.03.27.587047

**Authors:** Úrzula Franco-Enzástiga, Nikhil N. Inturi, Keerthana Natarajan, Juliet M. Mwirigi, Khadija Mazhar, Johannes C.M. Schlachetzki, Mark Schumacher, Theodore J. Price

## Abstract

Gene expression is influenced by chromatin architecture via controlled access of regulatory factors to DNA. To better understand gene regulation in the human dorsal root ganglion (hDRG) we used bulk and spatial transposase-accessible chromatin technology followed by sequencing (ATAC-seq). Using bulk ATAC-seq, we detected that in females diverse differentially accessible chromatin regions (DARs) mapped to the X chromosome and in males to autosomal genes. EGR1/3 and SP1/4 transcription factor binding motifs were abundant within DARs in females, and JUN, FOS and other AP-1 factors in males. To dissect the open chromatin profile in hDRG neurons, we used spatial ATAC-seq. The neuron cluster showed higher chromatin accessibility in GABAergic, glutamatergic, and interferon-related genes in females, and in Ca^2+^-signaling-related genes in males. Sex differences in transcription factor binding sites in neuron-proximal barcodes were consistent with the trends observed in bulk ATAC-seq data. We validated that *EGR1* expression is biased to female hDRG compared to male. Strikingly, *XIST*, the long-noncoding RNA responsible for X inactivation, hybridization signal was found to be highly dispersed in the female neuronal but not non-neuronal nuclei suggesting weak X inactivation in female hDRG neurons. Our findings point to baseline epigenomic sex differences in the hDRG that likely underlie divergent transcriptional responses that determine mechanistic sex differences in pain.

## Introduction

Women exhibit a higher prevalence of chronic pain [53], a greater number of clinical pain conditions [5], and a heightened sensitivity to experimental pain compared to men [24]. The strongest evidence is epidemiological and mechanistic sex differences in chronic pain conditions [45-47; 62]. Neuropathic, inflammatory and idiopathic chronic pain have higher prevalence rates, symptom severity and duration in women compared to men [16; 47; 78]. Although hormonal, immunological, anatomical and psychosocial factors contribute to sex differences in pain [55], at least some of these differences are likely driven by sex-specific gene expression in sensory neurons, glia or immune cells of the human dorsal root ganglion (hDRG). Support for this idea comes from animal model work demonstrating distinct mechanisms that promote and resolve chronic pain in rodents [20; 25]. Mechanistic studies in humans demonstrate sex differences in gene expression for transcription factors and epigenetic remodelers in hDRG from thoracic vertebrectomy patients suffering from chronic pain [50; 59]. Moreover, in both the human tibial nerve and in the human DRG, as well as in the mouse DRG, there are sex differences in gene expression in individuals without any pain history suggesting underlying epigenetic differences at baseline that cause these divergent patterns of gene expression [36; 42; 50; 58; 59; 75; 76].

Chromatin is a macromolecular complex composed of DNA, RNAs, and proteins found within the nuclear space. Epigenetic marks, such as specific histone or DNA modifications, can affect the degree of DNA compaction resulting either in open or accessible regions termed euchromatin or closed compact regions known as heterochromatin [3]. Chromatin accessibility is defined as the extent to which nuclear macromolecules can physically interact with chromatinized DNA [31]. Although chromatin serves a structural role by compacting DNA, it is also a dynamic entity that responds to cellular needs such as transcription, DNA repair, and differentiation [39; 44]. These dynamic changes in chromatin have been shown in the DRG in animal models in response to nerve injury and to inflammation, demonstrating a functional link between epigenomic regulation of gene expression and development and maintenance of chronic pain [9; 21; 34; 37; 41; 72; 80; 81]. To date very little is known about chromatin accessibility in hDRG although one previous study using microarray-based genomic and mRNA expression profiling found that expression quantitative trait loci were associated with open chromatin regions based on encyclopedia of DNA elements datasets [52]. A study of the human trigeminal ganglion (TG) found association between single nucleus ATAC-seq (snATAC-seq) open chromatin regions and single nucleotide polymorphisms (SNPs) associated with migraine [88]. However, the degree to which SNPs linked to migraine could be correlated to open chromatin in human TG cells was limited because snATAC-seq failed to cover all TG cell types detected with snRNA-seq.

We hypothesized that epigenomic differences in the hDRG may underlie mechanistic differences in molecular events that promote pain in females versus males and contribute to the higher propensity for women to develop chronic pain disorders. As a first test of this idea, we used both bulk and spatial ATAC-seq coupled with bulk nuclear RNA-seq to map the open chromatin accessibility landscape of DRG in female and male organ donor-recovered tissues. We found pervasive differences in transcriptional regulatory programs between females and males that likely have important implications for understanding epidemiological and mechanism-based differences in chronic pain in humans.

## Materials and methods

### DRG tissue preparation

All human tissue procurement procedures were approved by the Institutional Review Board at the University of Texas at Dallas. Lumbar DRGs from human organ donors were obtained via a collaborative effort with the Southwest Transplant Alliance. Right after dissection, human L4 or L5 DRGs from male and female organ donors were either transported in aCSF to be further processed within our facilities or frozen in dry ice right after dissection and stored in a −80 °C freezer. DRG donor demographic information is provided in **Table S1**. The inclusion criteria for this study took into account adult donors aged 18 to 65 years who exhibited no signs of pain or neuropathy. The cutoff age of 65 years old was determined based on prior epigenomic studies, which indicated significant alterations in both males and females occurring notably after the age of 65 [40].

### Bulk ATAC-seq on fresh hDRG

Immediately after dissection, human L4 or L5 DRGs from male and female organ donors were transported in cold bubbled NMDG-aCSF pH 7.4 (93 mM NMDG, 2.5 mM KCl, 1.25 mM NaH_2_PO_4_, 30 mM NaHCO_3_, 20 mM HEPES, 25 mM glucose, 5 mM ascorbic acid, 2 mM thiourea, 3 mM sodium pyruvate, 10 mM Mg_2_SO_4_, 0.5 mM CaCl_2_, 12 mM N-acetylcysteine; osmolarity 310 mOsm [82] in ice to our facilities. Using scissors, tissue was chopped in 2 mL of ice-cold nuclei isolation buffer [43]. This buffer contained: 250 mM sucrose, 25 mM KCl, 5 mM MgCl_2_, 10 mM Tris-HCl pH 8.0 and 0.1% triton X-100. The total volume was transferred to a Dounce homogenizer. 5 strokes of the loose pestle and 15 strokes of the tight pestle were applied. The homogenate was transferred to a conical tube and centrifuged at 100 x g for 8 min at 4 °C. The supernatant was carefully removed without disrupting the soft pellet. Next, the pellet was resuspended in 2 mL of nuclei isolation buffer free of triton. The supernatant was removed without disrupting the pellet after centrifugation at 100 x g for 8 min at 4 °C. Subsequently, the nuclei were resuspended in 2 mL of nuclei isolation buffer without triton and filtered through a 40-um cell strainer. The number of isolated nuclei was counted using a hematocytometer. A total of 100,000 nuclei were aliquoted and centrifuged at 500 x g for 10 min at 4 °C. The supernatant was then removed, and the pellet was utilized for the tagmentation reaction and DNA purification using the Active Motif kit (Cat. No. 53150). In brief, 50 uL of Tagmentation Master Mix (consisting of 2X Tagmentation buffer, 10X PBS, 0.5% digitonin, 10% tween 20, and assembled transposomes) was added to each sample. The tagmentation reaction was incubated at 37 °C for 30 min in a thermomixer at 800 rpm. Subsequently, 250 uL of purification binding buffer and 5 uL of 3 M sodium acetate were added. DNA purification columns were employed to isolate the DNA, and the tagmented DNA was eluted in 35 uL of DNA elution buffer. The amplification of tagmented DNA through PCR was carried out using i7- and i5-Illumina’s Nextera-based adapters as per the provider’s instructions. Pair-end 75 cycle sequencing reads were acquired on the Nextera500 sequencer or Nextera2000 sequencer. A total of 100 million reads were obtained from each sample.

### RNA-seq on fresh hDRG

After isolating 100,000 nuclei from hDRG for bulk ATAC-seq, the remaining extracted nuclei were used to perform RNA extraction with the aim of sequencing and pairing bulk nucRNA-seq with ATAC-seq from the same sample. Four male and 3 female hDRG samples were utilized to extract RNA, thus the pairing with ATAC-seq samples was performed in 6 out of 10 ATAC-seq samples.

Total RNA from hDRG nuclei was purified using TRIzol™. RNA peak profiles were analyzed using 5200 Fragment Analyzer and quantified with Qubit. RNA quality number was used to assess RNA Integrity (**Table S2**). Illumina Tru-seq stranded RNA library prep was used to generate cDNA libraries according to the manufacturer’s instructions. Next Generation Sequencing (RNA-seq) was performed in a NextSeq 2000 sequencer. Single-end sequencing was performed in a multiplexed fashion by loading the samples on a P2 flow cell, averaging ∼50 million reads per sample.

### Spatial ATAC-seq on frozen hDRG

The frozen hDRGs were gradually embedded in OCT in a cryomold by adding small volumes of OCT over dry ice to avoid thawing. A total of 8 hDRGs were used for these experiments (N=3 female, N=5 male) and demographic information of the organ donors is provided in **Table S3**. All tissues were cryostat sectioned at 10 μm onto SuperFrost Plus charged slides (Fisher Scientific; Cat 12–550-15). Sections were only briefly thawed in order to adhere to the slide but were immediately returned to the −20 °C cryostat chamber until completion of sectioning. The slides were removed from the cryostat and sent to AtlasXomics for further processing [13]. Spatial ATAC-seq (AXO-0303) consisted of the next steps: the tissue was fixed for 5 min with 0.2% paraformaldehyde. Afterwards, the tissue was treated with glycine and left to dry. Then, a tissue permeabilization step was performed using 0.1% NP40 for 15 minutes. Tn5 transposition on the fixed hDRG sections was carried out for 30 min. Adapters containing a ligation linker were added to the preparation to be inserted in accessible genomic loci. Barcodes with linkers were introduced using microchannels and were ligated to the 5′ end of the Tn5 oligo through successive rounds of ligation. hDRG sections were imaged to correlate spatially barcoded accessible chromatin with tissue morphology. A barcoded-tissue mosaic was created, reverse cross-linking was performed to release DNA-fragment, which were then amplified through PCR for subsequent library preparation. 150X150 paired-end sequencing was performed using NextSeq 2000 with 15% PhiX. Sequencing depth was up to 300 million reads per sample.

### Processing of bulk ATAC-seq data

First, we verified the quality of raw sequencing fastq files using FastQC v0.11.7. We removed adapters using TrimGalore-0.6.6 and verified the quality of trimming using FastQC v0.11.7. Paired-end reads were mapped to the reference genome GRCh38/hg38 using Bowtie2. To remove unmapped reads, low quality reads or reads that mapped to mitochondrial DNA and PCR duplicates we used Bamtools. Areas of open chromatin were identified by peak calling for each sample using Macs2 v.2.2.7.1 in keeping with Galaxy ATAC-seq data analysis. Differential accessibility between male and female donors was analyzed using DiffBind, a package within the Bioconductor framework. During differential analysis, normalization was performed in a default mode. Next, we used HOMER annotatePeaks which uses the program assignGenomeAnnotation to efficiently assign ATAC-seq peaks to genomic annotations. The definitions of annotations were acquired in a default mode. Processing of transcription factor binding sites was performed using Transcription factor Occupancy prediction By Investigation of ATAC-seq Signal (TOBIAS, [4]) in conjunction with human consensus transcription factor motifs from JASPAR. In brief, ATACorrect parameter was used to assess and correct the Tn5 transposase sequence preference of cutting sites. Putative transcription factor binding sites or transcription factor occupancy within peak regions were estimated using FootprintScore. Differential transcription factor binding sites in ATAC-seq reads between male and female hDRG were detected with BINDetect. Differentially Accessible Regions (DARs) were identified using DESeq2 v1.42.0.

### Processing of RNA-seq data

We confirmed the quality of raw sequencing fastq files using FastQC v0.11.7. The STAR alignment tool (v2.7.10b) was employed for aligning sequenced reads to the human GENCODE reference genome (v38). Sequenced reads underwent trimming to reduce sequencing quality issues at both ends. Relative abundance quantified as TPM was generated using the Stringtie tool and HTseq for counts generation.

### Processing of spatial ATAC-seq data

Processing of spatial ATAC-seq was performed by AtlasXomics, as previously reported [13]. Chromap was used to perform the alignment of sequenced reads to the human GENCODE reference genome (v38) and remove duplicates from fastq files [91]. Fragments files containing coordinates of sequenced DNA molecules mapping to barcodes were generated for downstream analysis. For data visualization, pixels on tissue samples were designated using AtlasXbrowser based on microscopy images to create Seurat-compatible metadata files. ArchR was used to process fragments files and the remotion of pixels not on tissue were removed. Data normalization and dimensionality reduction was applied to the data using ArchR’s iterative latent semantic indexing and subsequent graph clustering and Uniform Manifold Approximation and Projection (UMAP) embedding were performed. ArchR functions were used to acquire gene accessibility scores and matrices (gene score model), marker genes for clustering, and genome browser tracks. ShinyGO-080 was used to plot chromosomal location of DARs-associated genes to the human genome.

### *In situ* hybridization

hDRGs from non-pain donors were embedded in OCT using a cryomold over dry ice and sectioned. 10 µm sections for *XIST* and 20 µm sections for *EGR1* detection were placed onto SuperFrost® Plus charged slides (Thermo Fisher Scientific, cat. No. 1255015). Two sections per hDRG separated by at least 100 µm from each other were obtained to capture two sets of neurons allowing for a diverse population to be analyzed. Sections were fixed in cold 10% formalin pH 7.4 for 15 min and then dehydrated sequentially in 50% ethanol for 5 min, 70% ethanol for 5 min, and 100% ethanol for 10 min at RT. The slides were allowed to dry briefly and hydrophobic boundaries around the tissue sections were drawn with ImmEdge® PAP pen (Vector Laboratories cat. no. H-4000). RNAScope® Fluorescent Multiplex Reagent Kit (Advanced Cell Diagnostics (ACD), cat. no. 323100) was used to detect target *EGR1* (ACD, cat. no. 457671), *XIST* (ACD, cat. no. 311231) and *SCN10A* (ACD, cat. no. 406291-C2) mRNAs in hDRGs. The sections were treated with hydrogen peroxide (ACD, cat. no. 322335) for 10 min at RT followed by a rinse with ddH_2_O. Sections then underwent 5 s of protease III (ACD, cat. no. 322337) digestion followed by a quick wash in 1X PBS at RT. *EGR1* or *XIST* (Channel 1) and *SCN10A* (Channel 2) probes were mixed in 50:1 ratio. The probe solution was added to the sections placed onto the HybEZ™ Slide Rack (ACD, cat. no. 310017) which was then placed inside the HybEZ™ tray (ACD, cat. no. 310012). Probes were hybridized at 40°C for 2 h in HybEZ™ II Oven (ACD, cat. no. 321721). Sections were rinsed twice in 1X RNAScope® Wash Buffer (ACD, cat. no. 310091) at RT for a couple of minutes each. The sections were stored in 5X SSC buffer (Sigma-Aldrich, cat. no. S6636) overnight at RT. The slides were rinsed twice with 1X wash buffer followed by amplification with Amp 1 for 30 min, Amp 2 for 30 min, and Amp 3 for 15 min at 40°C. Channel 1 probe was coupled with Cy3 for *EGR1* or *XIST* detection and channel 2 with Cy5 for *SCN10A* detection. After rinsing two times with 1X wash buffer, sections were incubated with 1:5000 DAPI in 1X PBS for 1 min. Tissue sections underwent a rinse with 1X PBS before being air-dried completely and cover-slipped with ProLong™ Gold Antifade Mountant (Fisher Scientific cat. no. P36930). Slides were imaged at 20X, 60X and 100X magnification using the Olympus FV3000 RS confocal laser scanning microscope. In situ hybridization analysis was performed using ImageJ Fiji version 2.14.0 by Difference of Gaussians (DoG) edge detection method. For EGR1 analysis, ROIs were hand-drawn around each neuron avoiding lipofuscin on the image hyper stack. Channels with target probe were separated from the hyper stack and duplicated. The two images underwent the application of Gaussian Blur with sigma values 1 and 2 respectively. The two resulting images with blurs were subtracted using the built-in Image Calculator feature in ImageJ Fiji which allowed for detection of in situ hybridization puncta using the default threshold function. Thresholding values that allowed for as little noise as possible without compromising the signal were chosen for each channel. Analyze Particles feature was used to analyze the number of puncta. The count readout was extracted from the images across the 10 donors which were then used to plot *EGR1* puncta count in *SCN10A* positive or negative neurons for each sex. *EGR1* puncta count data are presented as violin plots with thick line indicating the median and the dotted lines indicating the quartiles. Data were analyzed with Graphpad Prism V9 (Graphpad, San Diego, CA). Student’s t test was used to assess differences in puncta count between sexes. A p<0.05 was considered statistically significant. Donor information for these experiments is provided in **Table S4**. For *XIST* analysis, ROIs were hand-drawn around analyzed nuclei using DAPI as a guide. Dispersal of *XIST* was measured based on dispersion of hybridization signal puncta in hand-draw ROIs that traced the analyzed nuclei. This area was divided by the total nuclear area stained with DAPI. Donor information for these experiments is provided in **Table S5**.

## Results

### Global chromatin landscape in female and male hDRG

To characterize the epigenetic landscape of the hDRG we performed ATAC-seq [8], an unbiased approach to assess chromatin accessibility regions within the genome [65] on lumbar hDRG from 9 donors (N=5 females, N=4 males, **Fig. 1A**). Donor information for all the samples that were used for bulk-ATAC in hDRG is shown in **Table S1**. After peak calling, we obtained a total of 669,749 peaks in female and 506,435 in male hDRG samples. At the global level, the distribution of called peaks in both female and male hDRG samples is concentrated in close proximity to the Transcription Start Site (TSS), with a notable accumulation of ATAC-seq signal directly at the TSS rather than in the immediate vicinity (**Fig. 1B**). The ratio between aggregate distribution of reads centered around the TSS (± 1 kb) and the signal flanking it is referred to as TSS enrichment (TSSe) score. TSSe is an important quality control parameter in ATAC-seq. We obtained a mean TSSe of 3.17 (**Fig. 1C**) similar to other reports using postmortem human tissue [10; 26]. Fragments of peaks in peaks (FRiP) defined as the measure of how many of the total reads are located within the peaks was greater than 0.13 in average. By mapping ATAC-seq peaks to genomic regions, we found that ∼15% of peaks mapped within the putative promoter, ∼40% were in distal intergenic regions, and ∼35% were found in introns (**Fig. 1D**). The enrichment of accessible chromatin in intergenic regions suggests that distal regulatory sequences, such as enhancers, non-coding regulatory elements in *cis*, can be accessed to regulate gene expression. In many cases, peaks located mainly in promoters [70], but also those harbored in the gene body (introns and exons) have been found to be good predictors of gene expression [23]. However, the proof of concept for the functionality of enhancers is tissue specific and requires further experimentation [77]. Since our results show that, at the global level, chromatin accessibility in hDRG showed similar annotation to called peaks between sexes, and the proportion was typical to other ATAC-seq reports [87], we aimed to conduct a sex-specific analysis identifying specific genes to investigate whether sex differences in pain could be associated with chromatin structure.

**Figure 1.**
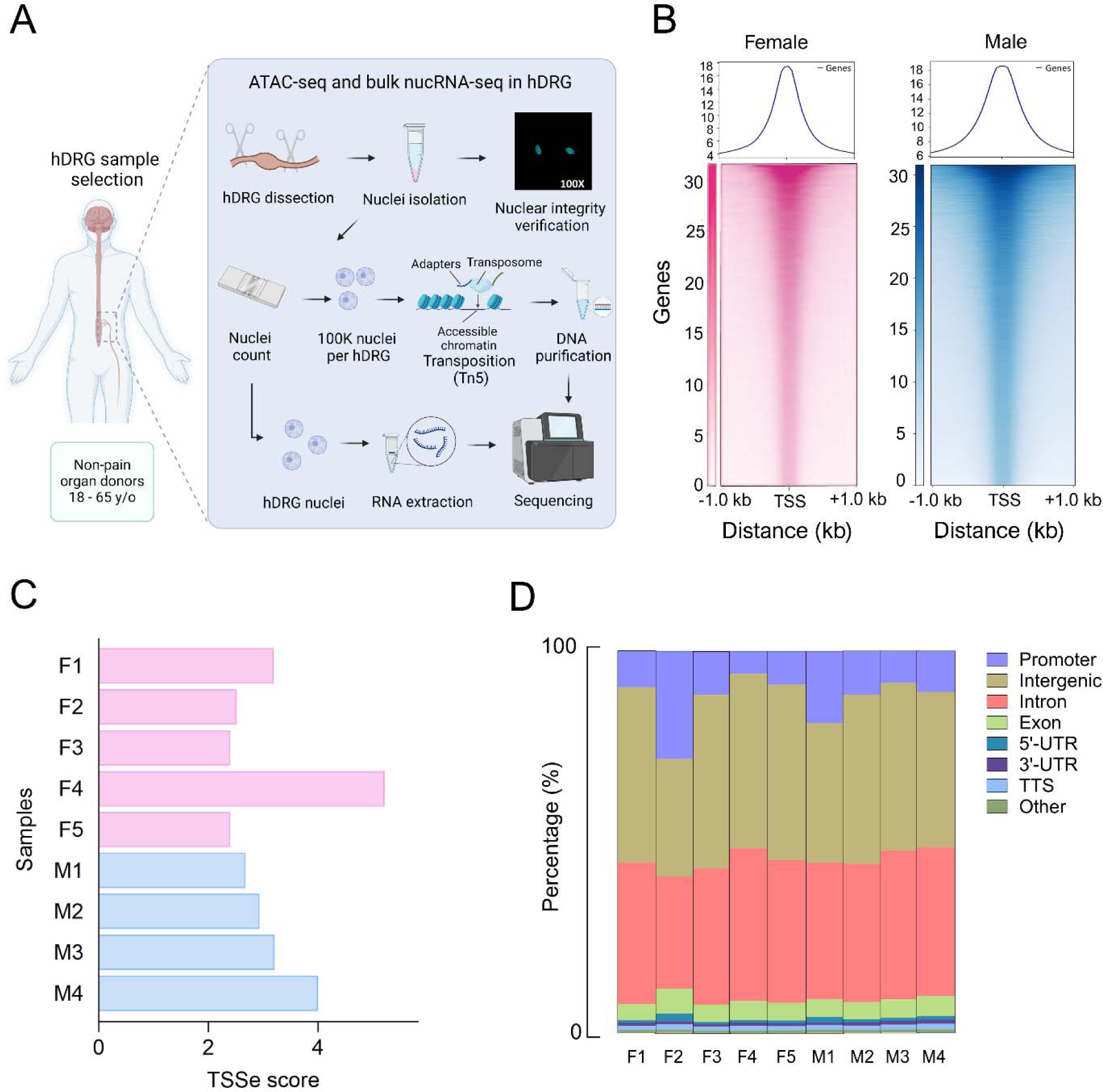
*Uncovering the open chromatin landscape in hDRG*. **A)** Schematic of the study design for bulk ATAC-seq. **B)** Heatmap shows chromatin accessibility around TSS ± 1 kb using aggregated peak enrichment scores from different female or male hDRG samples. Each row represents one consensus peak per gene spanning the TSS ordered by ATAC-seq signal intensity. Color represents the intensity of normalized chromatin accessibility. The metaplot on the top shows the mean ATAC-seq signal at the TSS. **C)** TSSe score obtained for each sample as a QC metric. **D)** Proportion of ATAC-seq peaks associated with different genomic annotations in each hDRG sample (UTR = untranslated region; TTS = Transcription Termination Site).

### Sex differences in chromatin accessibility in hDRG

By comparing ATAC-seq peaks in female and male samples, we found 3005 differentially accessible chromatin regions (DARs, FDR < 0.01, **Fig 2A**; complete DAR list is supplied in **Dataset S1**). Among these, 41 were identified in gene promoters in the female hDRG. Notably, the majority of DARs were located on the X chromosome in genes such as *MAPD72*, *FRMPD4, PORCN, PNMA3* and *TKTL1* (**Fig. 2A**). In males, 234 DARs were associated with gene promoters (±1kb from TSS) on autosomal chromosomes and included genes such as *NR1D1*, *IRF2*, *CCNB1*, *DAPK3* and *FOXO1* (**Fig. 2A**, promoter DAR list is supplied in **Dataset S2**). The ATAC-seq principal component analysis (**Fig. 2B**) demonstrated that female and male samples clustered separately, consistent with the DAR analysis results. Representative sex biased DARs around the TSS in female and male hDRG are shown in **Fig. 2C** and **D**, respectively. X inactivation is known to balance X chromosome gene expression between sexes, although it is incomplete in some tissues and new regulatory functions associated with escape from X inactivation are still being discovered [64]. We observed a striking number of DARs in this chromosome in the female DRG likely consistent with a lack of X silencing in the hDRG in females or the presence of poised genes ready to be activated. X inactivate-specific transcript (*XIST*), the long non-coding RNA (lncRNA) that primarily functions to silence one of the X chromosomes in each female cell, has been found to be dysregulated and participate in inflammatory and chronic pain in animal models [35; 68; 73; 83], suggesting a possible X inactivation dysregulation which could become more relevant in pathological conditions.

**Figure 2.**
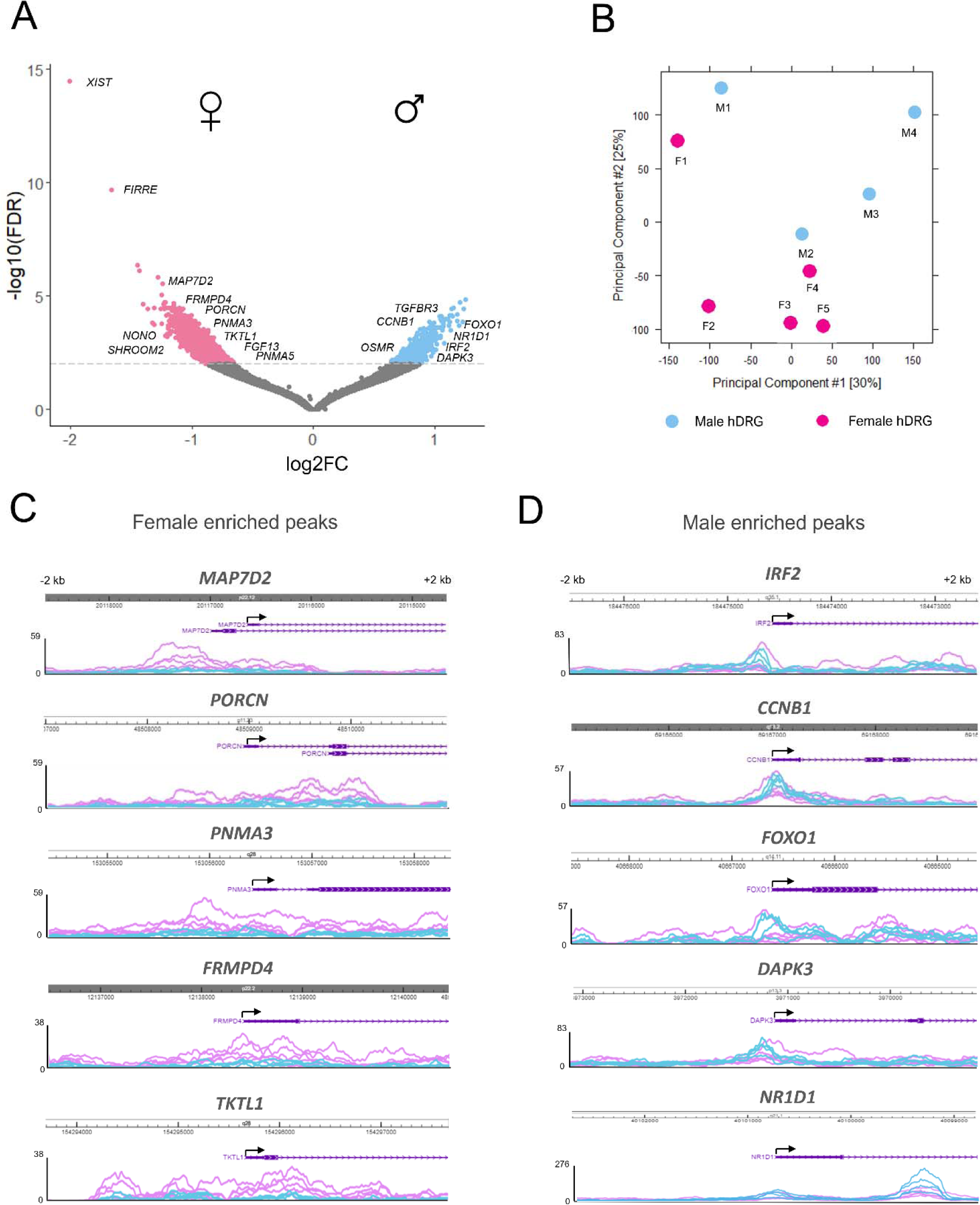
*Analysis of sex differences in open chromatin regions in hDRG*. **A)** Volcano plot showing DARs in hDRG between sexes (N = 9, 5 female (pink), and 4 male (blue) postmortem hDRGs, FDR < 0.01) identified with Diffbind. **B)** Principal component plot for ATAC-seq samples based on their similarity. **C)** Genome browser views of ATAC-seq signal of DARs surrounding the transcription start site from female hDRG. **D)** Representative genome browser tracks of DARs surrounding the transcription start site in male hDRG.

### Enriched transcription factor binding sites in sex-biased DARs in hDRG

In addition to chromatin accessibility, ATAC-seq data enables the prediction of transcription factor binding to open chromatin regions. With the aim of dissecting sex-specific transcriptional activity programs in hDRG, we analyzed the differential transcription factor binding motifs within accessible chromatin between sexes in hDRG using Transcription factor Occupancy prediction By Investigation of ATAC-seq Signal (TOBIAS). For this analysis, differential binding scores with - log10 (p-value) larger than the 95th percentiles (top 5% in each direction) were considered as significant [4]. In females, transcription factors of the EGR family such as EGR1 and EGR3, and members of the SP family such as SP1, SP2, SP4 and SP9 footprints were more abundant compared to male. In males, we found a higher level of footprints for transcription factors that are part of the activating protein 1 (AP-1) family such as JUN, FOS, FOSL2 and ATF4. (**Fig. 3A-B**). Binding of transcription factors to the promoter of specific accessible genes was also an output of TOBIAS analysis. We found that the EGR1 transcription factor has binding sites in *ADORA2B*, a synaptic-localized GPCR gene; *CX3CL1*, a chemokine known to participate in paclitaxel-induced macrophage recruitment in the DRG [27]; and *JAK1*, a kinase essential for cytokine and growth factors signal transduction. EGR3 showed open chromatin binding sites in the promoter of *PIEZO2*, a mechanosensitive ion channel known for contributing to sensory neuron functions [51]; *OXT, a* neuropeptide that suppresses action potential firing of DRG neurons [22; 57]; *GLRA3*, a subunit of the glycine receptor; and *TACR1*, a substance P receptor. The promoter of SP1 had binding sites on *TRPV2*, inflammation-related genes such as *TRAF1/4*, *STAT4* and *IKBKG*, and heat shock proteins like *HSF1*. Finally, SP4 showed open chromatin binding sites in Ca^2+^-signaling-related genes such as *CACNA1G* and *CARHSP1*, as well as the interferon regulatory transcription factor, *IRF6*. Examples of genes predicted to be regulated by binding of EGR and/or SP transcription factors are depicted in **Fig. 3C**. AP-1 transcription factor and JDP2 binding sites in males had diverse common targets in the accessible chromatin in hDRG, mainly in cytokine-related genes including *OSMR*, *IFI16*, *CCR7*, *NLRP3*, *TLR4* and *CASP8*. AP1 transcription factor binding sites were also found in GABAergic genes including *GABRA2* and *GABRB2* (**Fig. 3D**). Together, these results suggest that females and males have different transcriptional activation programs which could come play an important role in sex differences in response to injury.

**Figure 3.**
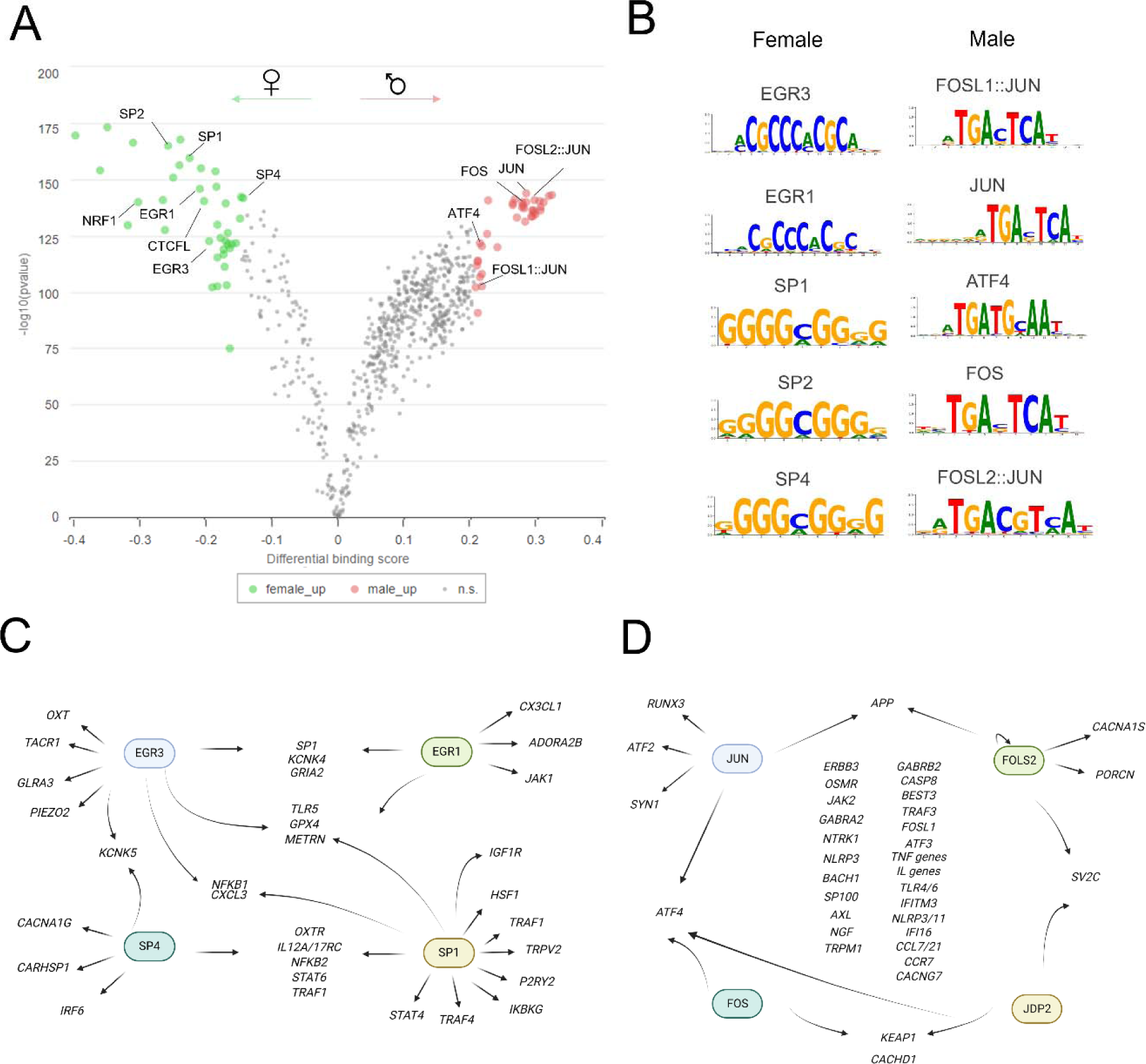
*Sex differences in transcription factor binding motifs within ATAC-seq peaks.* **A)** Volcano plot showing the differential transcription factor footprints enriched in female and male hDRG identified with BINDetect (N = 9, 5 female and 4 male postmortem hDRGs, -log10 (p-value)> 95th percentiles). **B)** Differential transcription factor consensus footprints in female and male hDRG defined by reference motif on JASPAR repository. **C**) Genes predicted to bind EGR/SP factors due to transcription factor footprints in their promoters determined with TOBIAS. **D**) Genes with transcription factor binding sites for AP-1 factors in their promoters identified with TOBIAS.

### Spatial ATAC-seq in hDRG

Since we did not perform neuronal selection for our bulk ATAC-seq experiments in hDRG, and spatial information is lost during dissociation, we used spatial ATAC-seq to dissect the chromatin profile in neuronal barcodes in a similar fashion to how we have done previously in spatial RNA-seq [76]. Spatial ATAC-seq was performed in hDRG tissue sections from 8 donors (N = 5 male, N =3 female) using a microfluidic barcoding system, followed by next generation sequencing [13].

We obtained 45,836 fragments on average in hDRG tissue, a TSSe score of 5.426 and a FRiP of 0.1276 of fragments of reads in peaks (**Table S6**). The improved TSS score may be due to tissue preparation, since DRGs for spatial experiments were immediately frozen in the operating room while DRGs for bulk ATAC-seq experiments were transported for 30-60 min in cold artificial cerebral spinal fluid (aCSF) prior to nuclear isolation and subsequent tagmentation (see **methods**). The average proportion of mitochondrial fragments was 4.15% (**Table S7**), and these reads were removed from the analysis. ATAC-seq fragment size distribution showed a typical periodicity of ∼200 bp corresponding to reads, consistent with the length of DNA protected by mono-, di-nucleosomes (200, or 400 bp), as well as nucleosome free DNA (<100 bp) or open chromatin regions (**Fig. S1**), all of which is consistent with typical nucleosome packing [7].

In addition to neurons, there are at least 8 broadly defined cell types in hDRG [6; 30; 49]. This high hDRG cellular heterogeneity was also observed at the epigenomic level, since unsupervised cell clustering in spatial ATAC-seq samples identified 9 groups of cells (**Fig. 4A**). Cluster 1 was classified as a neuronal cluster due to the abundance of chromatin accessibility in *SNAP25* gene compared to other cell types. Further, this cluster also shows chromatin activity for *CALCA* and *SCN10A* consistent with a neuronal phenotype (**Fig. 4B-D**). Cluster 2 showed enriched chromatin accessibility for the marker of satellite glial cells (SGCs), *FABP7* (**Fig. 4E**). The chromatin accessibility profile for cluster 3 had *CD300LB, IFI30*, *C5AR1*, *C1QB* and *ABI3* among the top 30 most active genes, all of which are expressed by immune cells in the hDRG ([6] **Fig. 4F**). According to the hDRG harmonized atlas [6], the top 30 most active genes for the cluster 5 exhibited enrichment for immune-related genes, such as *OLR1*, *MS4A7*, *TYROBP* and *CORO1A*. Interestingly, this cluster also showed enrichment for macrophage markers such as *CD163, ARHGAP30, SIRPB2* and *PLCB2*. Thus, we classified cluster 5 as immune macrophage enriched (**Fig. 4G**). Cluster 6 showed enriched chromatin accessibility in *SCN7A* likely representing the non-myelinating Schwann cell cluster (**Fig. 4H**). Cluster 7 showed enriched chromatin accessibility in *PRX* and *MPZ* representing the myelinating Schwann cell cluster (**Fig. 4I**) A fibroblast cluster was identified by the gene activity of *DCN* and *FMOD* in cluster 8 (**Fig. 4J**). Cluster 9 was identified as the endothelial cluster due to abundant *CLDN5* gene accessibility (**Fig. 4K**). Cluster 4 was more challenging to classify. This cluster showed chromatin accessibility in genes such as *PODN* and *FOXD1*, all of which are expressed in fibroblasts in hDRG suggesting this cluster is composed of a subtype of fibroblast in the hDRG. Cell type composition in hDRG section shows consistent cluster proportion between samples and sexes (**Fig. 4L**). The number of unique fragments per barcode exceeded the minimum values required to be considered high quality in all the clusters (log_10_ nFrag = 3, **Fig. 4M**). TSSe scores values per cluster and nucleosome ratio are depicted in **Fig. S2**. The number of final peaks identified across all donors was 303,850 and the genomic annotation proportion was higher in intron and distal peaks, similar to our bulk ATAC-seq experiments (**Fig. 4N**).

**Figure 4.**
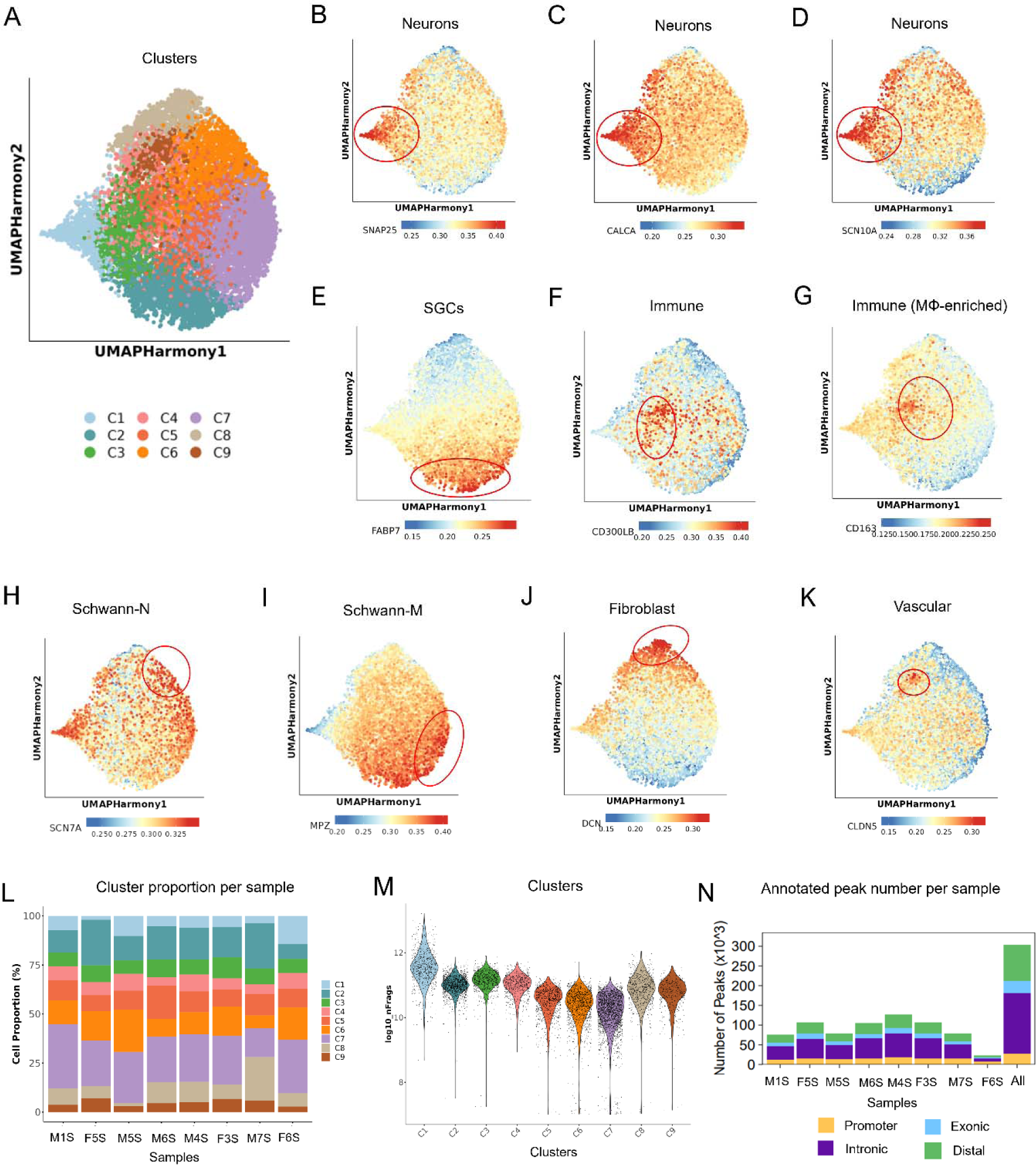
*Characteristics of spatial ATAC-seq in hDRG*. **A**) UMAP plot showing the 9 clusters generated by Seurat’s workflow. **B-D**) UMAP plots of the chromatin accessibility of gene markers that were used to label the neuronal cluster. **E-K**) UMAP plot showing cellular populations other than neurons, including satellite glial cells (**E**), immune cells (**F**), immune cells enriched in macrophages (**G**), Schwann-N cells (**H**), Schwann-M cells (**I**), fibroblasts (**J**) and vascular cells (**K**). **L**) Consistent cluster proportion in each sample. **M**) Violin plot shows number of unique fragments per cluster. **N**) Number of peaks identified across all donors (all), and within samples associated and their genomic annotations: promoter, intron, exon and intergenic (N = 3 female, N =5 male; samples M5S, M6S, M7S and F6S belonged to donors whose samples had not been used before for bulk ATAC-seq. A frozen hDRG sample from donors M1, M4, F3 and F5 were used for spatial ATAC-seq, and are labeled as M1S, M4S, F3S and F5S).

### Pseudo-bulk analysis of spatial ATAC-seq data

We conducted a pseudo-bulk differential analysis between sexes using the reads from all the clusters in spatial ATAC-seq to have a general overview of sex differences in hDRG chromatin accessibility, which we hypothesized would be similar to our bulk ATAC-seq data (**Fig. 5**). In **figure 5A** we show that the number of fragments per barcode and the distribution of reads in the UMAP were similar between female and male hDRG samples (**Fig. 5B**). Our analysis showed DARs in 3796 genes in female hDRG versus 3827 genes in males (**Fig. 5C**). In females, a significant proportion of DARs was located on the X chromosome consistent with our bulk ATAC-seq findings (**Fig. 5D**). These findings align with our bulk ATAC-seq results. Genes such as *NONO*, *MAPD7*, *FRMPD4*, *PORCN*, *FGF13* and *TKTL1* shown in female bulk ATAC-seq were also found to be differentially accessible in female spatial ATAC-seq data. Spatial ATAC-seq also revealed female DARs in genes outside the X chromosome such as *ERBB4*, GABAergic genes including *GABRG1/2*, *GABRA5*, *GABRB3*, *GABRB1*, *GABRA2/4*, and other neuronal genes such as *BDNF*, *RELN*, *SCN9A*, *GRIA4* and *GRIN2B* (**Dataset S3**). It is important to mention that most of these genes are also significantly differentially accessible when only autosomes are analyzed, which validates our results (data not shown).

**Fig. 5.**
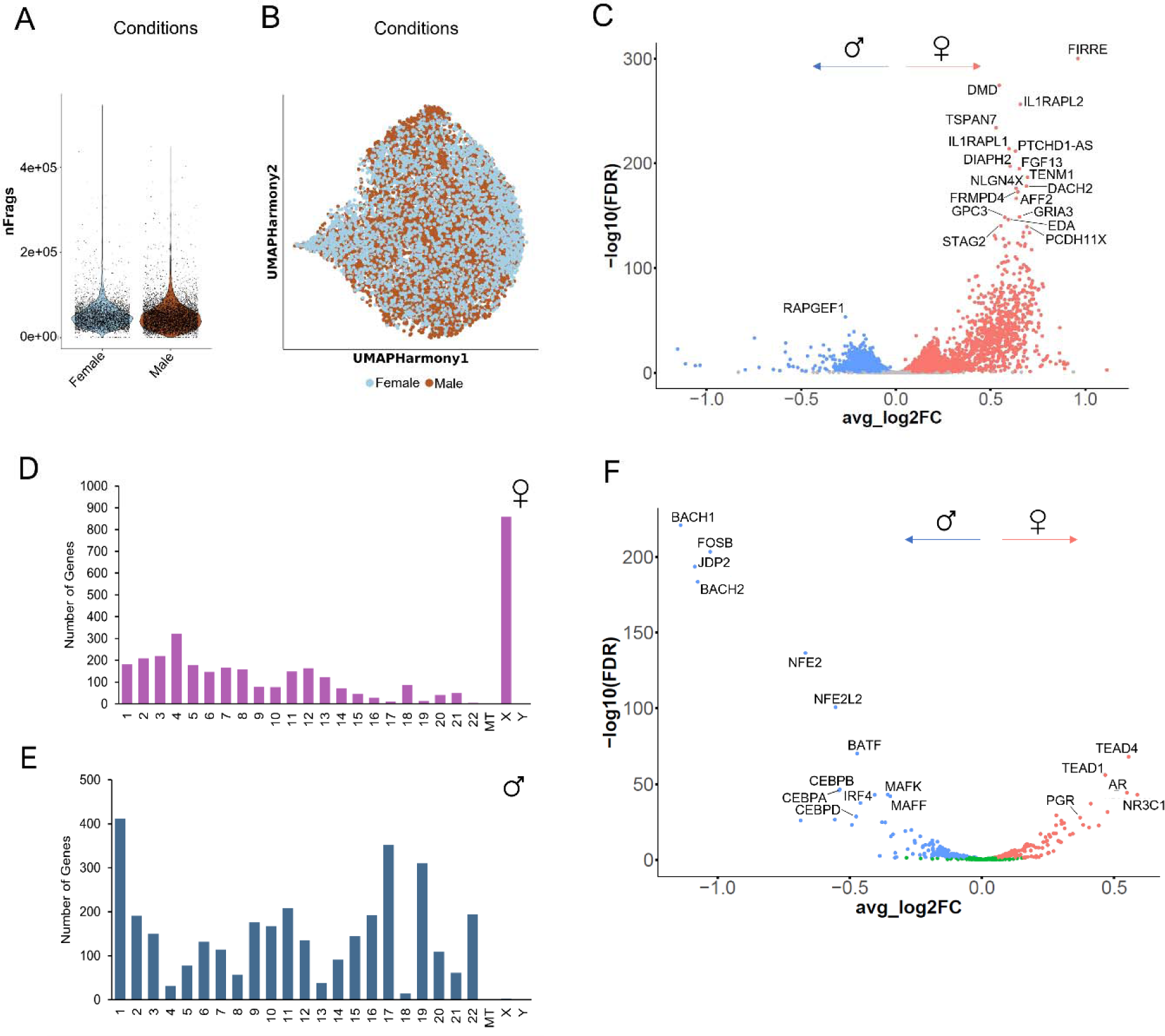
*Differential spatial ATAC-seq signal in hDRG between sexes*. Analysis of sex differences in chromatin gene activity in hDRG. **A**) Number of fragments per barcode in female and male hDRG samples. **B**) UMAP of female and male barcodes distribution along hDRG cellular populations. **C**) Volcano plot showing differential ATAC-seq gene activity in a pseudo-bulk fashion in hDRG in female vs male (N = 4 female, 5 male postmortem hDRGs), FDR < 0.01 identified with ArchR. **D**) Genome location of DARs in female spatial ATAC-seq spanning the human chromosomes generated with ShinyGO. **E**) Genomic position of DARs in male hDRG. **F**) Volcano plot showing differential transcription factor binding sites in female vs male hDRG (N = 4 female, 5 male postmortem hDRGs), FDR < 0.01 identified with ArchR.

On the other hand, in males, the peaks spanned the whole genome, with a higher abundance of DARs in genes located on chromosomes 17 and 19 (**Fig. 5E**). Furthermore, a considerable amount of the genes with DARs in promoters in our bulk ATAC-seq data were also found to be biased towards males in the spatial ATAC-seq. We found male hDRG DAR-associated genes such as *FOS*, *FOSL2*, *NR1D1*, *IRF2* and *FOXO1*. Male hDRG DAR list also included genes encoding for cation channels such as *TRPV2/4*. Interestingly, the male hDRG DAR list also included circadian related genes such as *BHLHE40* and *CSNK1D,* cytokines such as *CSF1R*, *CCL2*, *IL4/6R*, and alarmins *HMGA1*. Hormonal associated genes such as *INSR* and *THRA* were also found to be epigenetically more active in males. Finally, genes encoding for epigenetic remodelers such as *DOT1L*, *HDAC4*, *JARID2* and *ARID5B* were found in male hDRG.

Transcription factor (TF) motif enrichment analysis allowed us to infer differential binding sites in spatial ATAC-seq data. In females, transcription factors such as TEAD1/4, NR3C1, AR and PGR were enriched in the spatial ATAC-seq dataset. In males, peaks in DARs again belonged to the AP-1 family such as FOSB, or AP-1 interacting factor like BACH1/2 (**Fig. 5F**).

### Sex differences in chromatin accessibility in hDRG neurons determined with spatial ATAC-seq

A primary motivation of our spatial ATAC-seq approach was to identify potential sex differences in neurons in hDRG. To examine this, we compared chromatin accessibility in the neuronal-enriched cluster between sexes (**Fig. 6**). DESeq2 analysis revealed DARs associated with 1665 genes in females and 1089 genes in males (p_adj_ < 0.01, **Fig. 6A, Dataset S4**). Similar to what we observed previously with bulk ATAC-seq or the pseudo-bulk analysis of spatial ATAC-seq data, most of the peaks in females were located on the X chromosome (p_adj_ < 0.01, 643 out of 1665 genes, **Fig. 6B**). In male neurons, the genes were distributed across the entire genome, and again, there was a higher abundance observed on chromosome 17 and 19 (**Fig. 6C**).

**Fig. 6.**
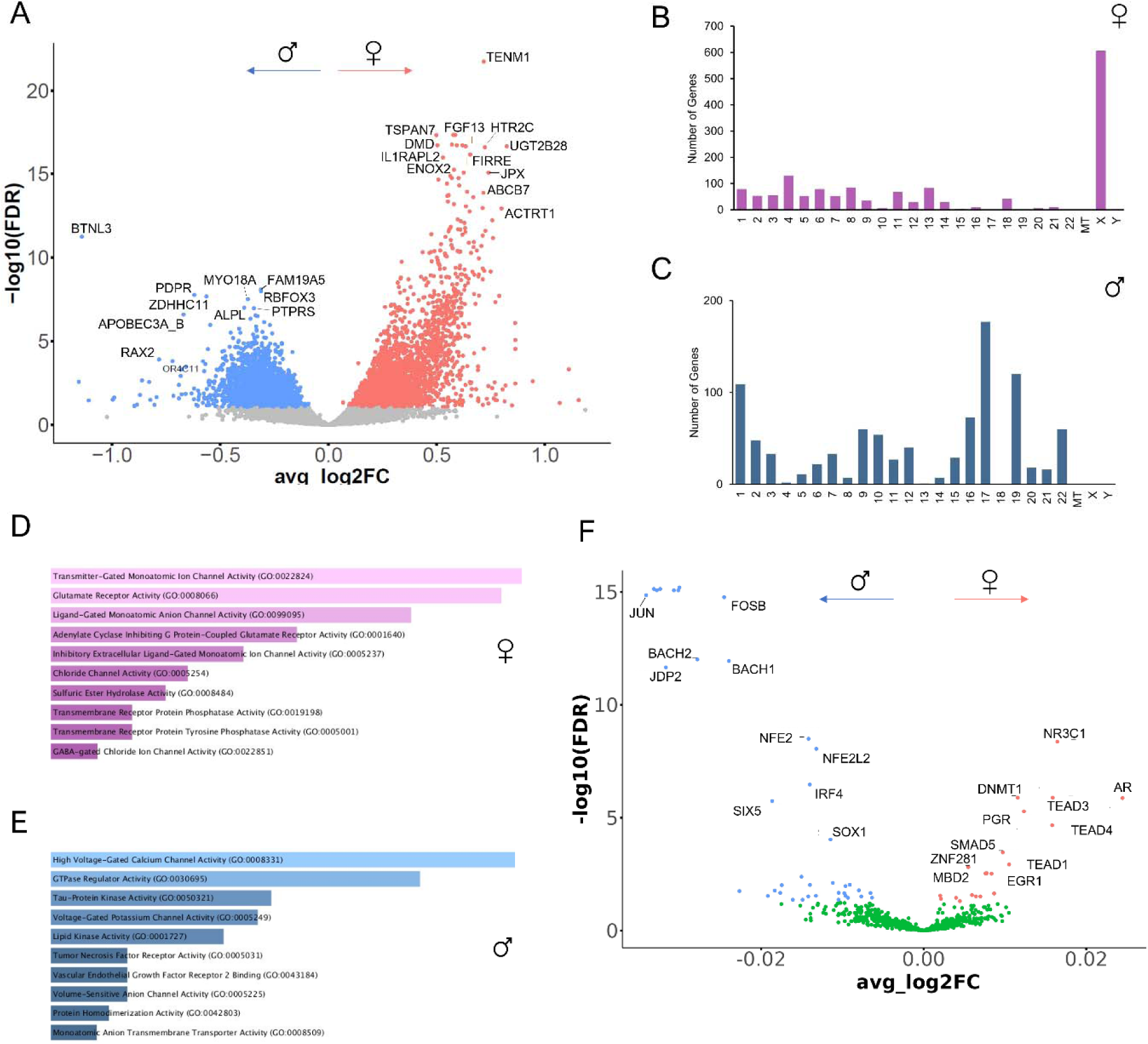
*Sex differences in spatial ATAC-seq DARs in the neuronal cluster*. **A**) Volcano plot showing DARs in spatial ATAC-seq neuronal clusters in hDRG neurons in female vs male (N = 4 female, 5 male postmortem hDRGs, FDR < 0.01 identified with ArchR. **B**) Genome location of DARs in female neuronal ATAC-seq spanning the human chromosomes generated with ShinyGO. **C**) Genomic position of the DAR gene set in males. **D**) Overview of the molecular function gene ontology for DAR-associated genes in female neurons in hDRG. **E**) Summary of the gene ontology related to molecular functions for DAR-associated genes in male neurons in hDRG. **F**) Volcano plot showing transcription factor binding site enrichment in DARs in hDRG neurons in females vs males (N = 4 female, 5 male hDRGs), FDR < 0.01 identified with ArchR.

The analysis of gene ontology of genes associated to DARs between sexes showed that glutamatergic genes such as *GRIA2* and *GRIA4* to be found in the top 100 DARs in female hDRG chromatin. GABAergic genes such as *GABRB1*, *GABRG1*, *GABRE*, *GABRQ*, *GABRA1/2/3/6* were also DARs in female hDRG within the differential female gene-set, together with the lncRNA *BDNF*-*AS*. Enrichment analysis of accessible regions showed an association between open chromatin regions and glutamate receptor activity as the main gene ontology enriched molecular function category (pval: 0.00001581, **Fig. 6D**) determined with EnrichR [32]. Interestingly, neurons in female donors also showed DARs in *ACE2*, some interferon related genes such as *IFNB1*, *IFNA5*, *IFIH1*, *IFI44* and *CASP4*, and centromere genes such as *CENPE/W* compared to males.

In male neurons, genes such as *GRIK3* and *PTPRS* were found among DARs compared to female. Further, voltage-gated Ca^2+^ channel related genes such as *CACNA1G, CACNA1I* and *CACNA2D2* were found in the male DAR gene set. Interestingly, *TRPV3* was also a DAR in males compared to female neurons. Gene ontology analysis of the male DAR gene set revealed high abundance of calcium channel activity in the molecular function category (p_val_ = 0.00165, **Fig. 6E**).

Finally, we analyzed the neuronal cluster for enrichment of transcription factor binding sites in the DARs between females and males. In females, *EGR1*, *TEAD1/3/4*, *NR3C1, AR* and *PGR* were enriched, and factors of the AP-1 family including *JUN* and *FOSB* as well as, *BACH1/2* and *IRF4* were found enriched in males (**Fig. 6F**).

### *EGR1* expression in hDRG

One of the outcomes that showed consistency in bulk and spatial ATAC-seq was the bias toward female in the footprint scores for the transcription factor EGR1 in hDRG, and specifically in neurons. With the aim of validating this consistent finding we sought to assess the expression of *EGR1* in hDRG using *in situ* hybridization. We found that *EGR1* is expressed in female and male hDRG *SCN10A* (+) and (-) neurons, but also in cells surrounding neurons, likely satellite glial or immune cells (**Fig. 7A-B**). We quantified the number of *EGR1* mRNAs localized to hDRG neurons and found it to be more abundant in female compared to male (**Fig. 7C**). This sexual dimorphism was statistically significant in *SCN10A* (+), but not *SCN10A* (-) cells in females (**Fig. 7D-E**). Our analysis also showed that females have a higher number of *EGR1* (+) *SCN10A* (+) cells compared to males (57.9% vs 29.71%, respectively, **Fig. 7F**). These results are consistent with our data sets. The combined identification of *EGR1* female-biased expression and the prominent abundance of regulatory binding sites in female hDRG, renders the transcriptional activity of *EGR1* a strong candidate for a female-biased regulator of gene expression in hDRG neurons.

**Fig. 7.**
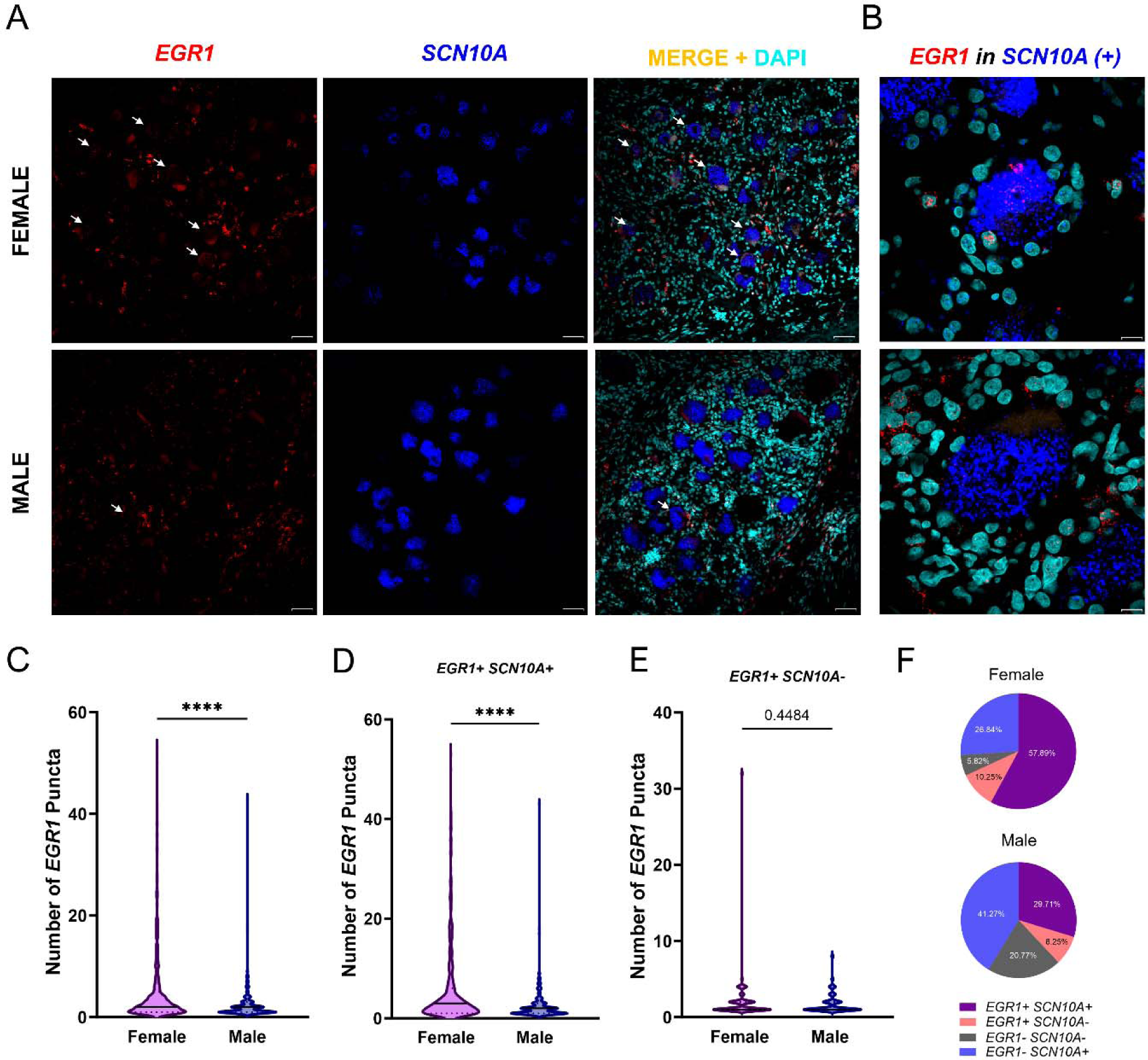
*EGR1* is expressed in more *SCN10A* positive neurons in females compared to males in hDRG. **A)**. Representative *in situ* hybridization images of *EGR1* (red) in *SCN10A* (+) neurons (blue) in female and male hDRG. *EGR1* positive neurons are indicated with an arrow. **B**) 100 X magnification of neurons with *EGR1* mRNAs in female and male neurons. **C**) Total number of *EGR1* mRNAs in neurons. **D-E)** Number of *EGR1* mRNAs in *SCN10A* positive and negative neurons, respectively. **F)** Percentage of *EGR1* positive cells in *SCN10A* positive or negative cells across female and male donors in hDRG. A total of 722 neurons were analyzed to assess *EGR1* expression in female hDRG, and 727 in males (N = 5 female, N = 5 male DRG). Data are presented as violin plots with thick line indicating the median and the dotted lines indicating the quartiles. Section thickness – 20 µm. Scale bar – 50 µm for **A** and 10 µm for **B**. ****p<0.0001 as determined by t test in **C-E**.

### Patterns of chromatin accessibility in hDRG are moderately positively correlated to the transcriptomic profile

In light of the sex dimorphic differences in chromatin accessibility between sexes in our bulk ATAC-seq results, we conducted a differential gene expression analysis on bulk nuclear RNA-seq (nucRNA-seq). RNA extractions and sequencing was performed in nuclei extracted from DRGs of 7 donors (N=3 females, N=4 males (same set of donors as in bulk ATAC-seq; details in **Table S1**). We found 8 genes that were statistically different between sexes in the nuclear compartment (p_adj_<0.05, fold change >1.33, **Dataset S5**). Females showed higher expression of chemokine genes such as *CCL3* and *CCL4*. As expected, we found *XIST* to be differentially expressed compared to male. An interesting point to highlight is that *EGR1* and *EGR3* were trending to be highly expressed in females compared to males in our RNA-seq data (**Fig. S3A**), which aligns well with transcription factor footprints in ATAC-seq data sets and in situ hybridization experiments. On the other hand, males showed higher gene expression of non-coding genes such as *KDM5DP1*.

We tested the correlation between the log2 fold change in DARs situated in promoter and within gene bodies [23], including exon, and intron regions in ATAC-seq and the log2 fold change in nucRNA-seq for those genes in the matched hDRG samples. We found a weak positive correlation between the transcriptional and the chromatin accessibility profiling (**Fig. S3B**). This weak correlation could be explained by the repertoire of RNAs analyzed, which were limited to nuclear space and not to the whole cell. Evidence has shown that nuclear compartment harbors abundant lncRNAs and small nucleolar RNAs in human brain tissue, consistent with the expressed genes in our nucRNA-seq data, while the cytoplasm contains RNAs associated to cellular function such as metabolism, translation and protein organization compared to the nuclear compartment [90]. In addition to the small sample size, another explanation is that a high correlation was not observed since accessible regions are not necessarily associated with actively transcribed genes, or they are associated with genes that are poised to become active under specific conditions [70]. It is important to denote that transitions in the deposition of histone modifications as well as transcription factor binding to regulatory sites such as enhancers can precede gene expression changes, inducing poised or primed chromatin states [28], which are indistinguishable from active chromatin in ATAC-seq due to a lack of epigenomic context. Further research combining different histone and DNA modifications is required to better understand the epigenomic landscape and to get a better interpretation of gene regulation in this complex tissue.

### *XIST* nuclear dispersion in female hDRG neurons

Both ATAC-seq datasets showed high accessibility for many X chromosome genes selectively in females, suggesting incomplete inactivation of the X chromosome in human, female DRG cells [1]. We explored the expression and localization of XIST RNA in hDRG because the RNA controls X inactivation to achieve gene dosage compensation between sexes by coating and inactivating one X chromosome in females [56; 61]. Although *XIST* is generally considered an exclusive long non-coding RNA in females, *XIST* has been shown to be lowly expressed in male nerve tissue [63]. We performed *in situ* hybridization for *XIST* in female and male hDRG. In females, *XIST* was expressed in neuronal and non-neuronal cells (**Fig. 8A**). Strikingly, we observed that *XIST* signal was not confined to a single area inside the neuronal nuclei. Instead, *XIST* signal was dispersed covering most of the nuclear compartment (**Fig. 8B**). This phenomenon was observed in *SCN10A*(+) and *SCN10A*(-) cells, where X spanned around 80% of the nuclear space. This observation was different in the cells surrounding the neurons, where *XIST* signal was more confined, being restricted to around 20% of the nuclear space (**Fig. 8C**). Compared to females, *XIST* was barely detected in male hDRG (**Fig. S4**). Since dispersion of XIST RNA in the nucleus is associated with incomplete X inactivation [74], these finds support the conclusion that hDRG neurons in females have incomplete X inactivation, consistent with our ATAC-seq findings.

**Fig. 8.**
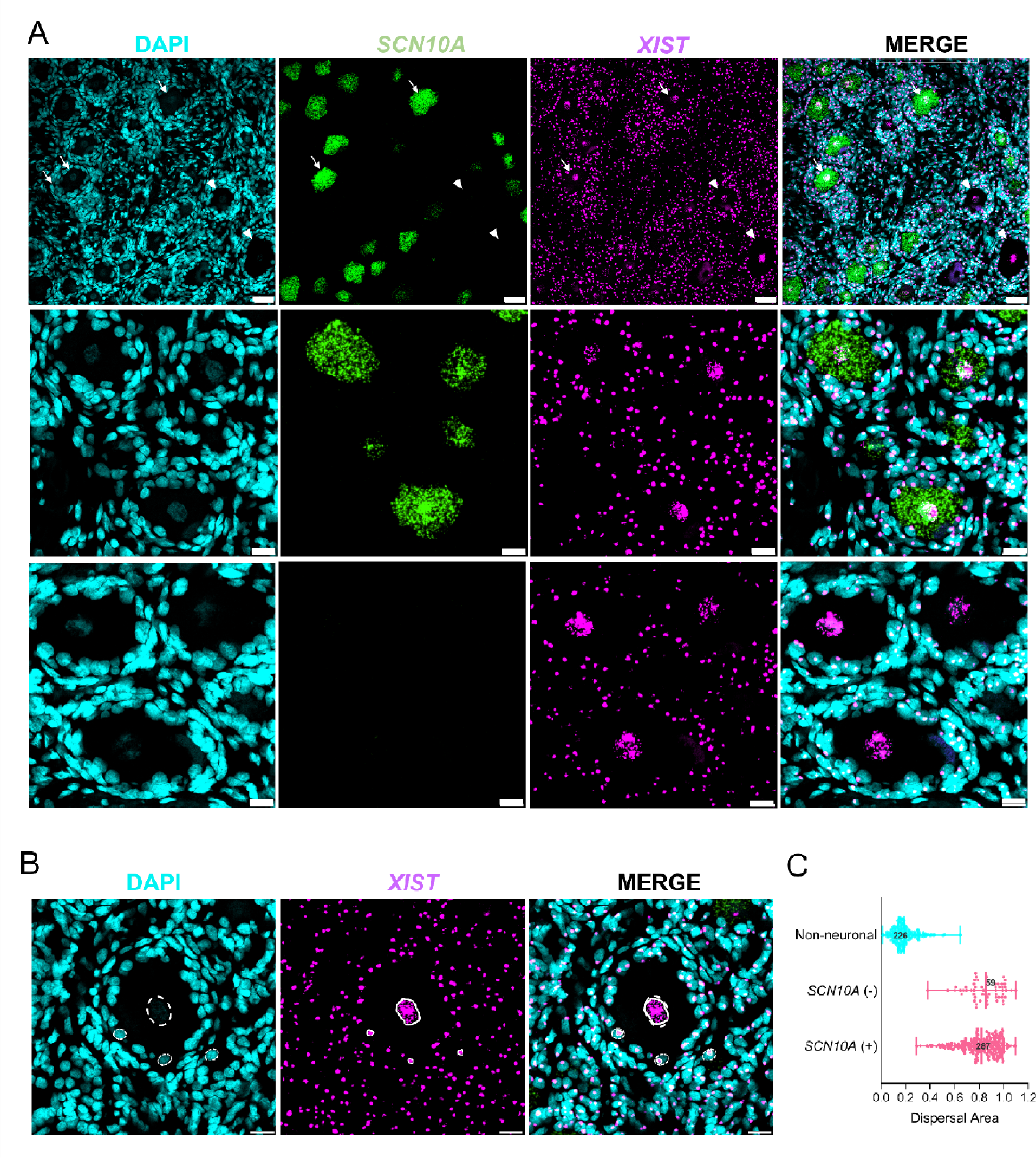
*XIST* is highly dispersed in the nuclear compartment of female neurons in hDRG. **A)**. Representative *in situ* hybridization images of *XIST* (pink) in *SCN10A* (+) and *SCN10A* (-) neurons (green) in female hDRG (20X images (top), and 3X zoom magnification (bottom)). *XIST* positive neurons, also positive for *SCN10A*, are indicated with an arrow. *XIST* positive neurons, negative for *SCN10A*, are indicated with an arrowhead. **B**) Representation of *XIST* dispersal analysis; an ROI was drawn around the nuclear compartment of neurons and non-neuronal cells (white). To delineate XIST area dispersion we drew an ROI with the dots that were furthest apart from each other (yellow). **C**) Representation of *XIST* dispersal area in *SCN10A* positive and negative neurons, and non-neuronal cells. A total of 346 neurons were analyzed to assess *XIST* expression and 226 non-neuronal cells in in female hDRG (N = 3 female, N = 2 male hDRG). Data are presented as scatter plots with lines indicating the mean and quartiles. Section thickness – 10 µm. Scale bar – 50 µm for **A** top and 20 µm for **A** bottom, and 20 µm for **B**.

## Discussion

Our work provides a comprehensive view of the chromatin accessibility profile in hDRG and focuses on sex differences in tissues recovered from organ donors without any history of chronic pain or neuropathy. Chromatin structure plays a crucial role in transcription through the controlled access of regulatory factors to DNA. We mapped the accessible DNA in hDRG using two different approaches, bulk and spatial ATAC-seq. The latter approach resolved hDRG chromatin accessible profiles in distinct cell types where we focused on neuronal signatures. First, our work shows a sexual dimorphism at the chromatin level between sexes in hDRG. This dimorphism is largely influenced by the X chromosome in female hDRG suggesting a reduced X inactivation or a dormant transcriptional state that can readily become activated under certain circumstances. Second, our ATAC-seq data show a sex dimorphic signature in transcription factor bindings sites in accessible chromatin. EGR and SP motifs were more prominent in female hDRG, while AP-1 and IRF were more abundant in males. Finally, our work shows sex differences in neuronal chromatin using spatial ATAC-seq. In neurons, sex differences were partly driven by the X chromosome in females but using this approach we also uncovered differences in autosomal genes. GABA-A channels and glutamatergic genes were the molecular ontologies associated with accessible DNA in females, while voltage gated Ca^2+^ and K^+^ channel genes as well as GTP associated proteins were associated to male hDRG open chromatin.

Our bulk ATAC-seq findings point to a high chromatin accessibility in the X chromosome in females. Independently, spatial ATAC-seq confirmed this finding. It is known that one of the two X chromosomes in females is epigenetically silenced to compensate for the imbalance of X dosage between sexes [56]. X inactivation is triggered by the upregulation of *XIST* early in development. XIST coats one of the X chromosomes in *cis* [61; 66] inducing X heterochromatization via repressive epigenetic signatures [54]. Interestingly, a small subgroup of genes escapes inactivation and have epigenetic marks of active transcription, allowing gene expression from the inactive X (Xi) chromosome [1]. Our findings are consistent with previous observations showing tissue-specific varying levels of X inactivation in human tissues [79]. Further, we showed that *XIST* is highly expressed in females, and its chromatin accessibility is higher in the adult female DRG. In situ hybridization experiments show concordant results, with female having a higher abundance of *XIST* in hDRG, which is barely detected in males. Strikingly, *XIST* was found highly dispersed in the neuronal nuclei but not in the nuclei of surrounding cells in hDRG. This phenomenon has been observed in human pluripotent cells [14], where it has been hypothesized that *XIST* regulates autosomal genes in females. Future experiments should test the idea of a reduced coverage of XIST on the Xi and the consequence of a high *XIST* nuclear dispersal in hDRG neurons. Our results are also consistent with a recent study showing greater variability in X inactivation in excitatory neurons in the human brain suggesting enhanced expression of genes from this chromosome in women in this population of neurons that shares a glutamatergic biochemical phenotype with DRG neurons [84].

It is now recognized that sex dimorphisms in pathological conditions like autoimmune diseases, can be driven by tissue or cell-specific patterns of X-inactivation. We observed higher chromatin accessibility mainly in genes coding for GABA-A receptor subunits, glutamatergic and interferon-related genes in females, some of which are located in X chromosome. This is in line with previous findings showing that interferon-related genes are more highly expressed in female hDRG from neuropathic pain patients undergoing thoracic vertebrectomy surgery [59]. *ACE2,* an interferon stimulated gene located in the X chromosome, is expressed in hDRG neurons, serving as potential entry site for SARS-CoV-2 in the PNS [69]. Delving deeper into the study of X inactivation escape in hDRG will shed light on the organization of chromatin in this chromosome and improve our understanding of its implications for chronic pain mechanisms in women. Importantly, a previous study linked X inactivation escape in blood cells to development of chronic pain after motor vehicle accident supporting the relevance of this line of investigation [89]. On the other hand, some of the female-biased genes were associated with DARs on autosomes, such as *GABRA5*. It has been shown that the α_5_-GABA_A_ receptor (encoded by the *Gabra5* gene in mice) is more highly expressed in the female rodent DRG where it plays an important role in pain signaling and is epigenetically regulated in a sex-specific fashion [17; 18]. We also found a higher accessibility in autosomal *BDNF*-*AS*, a lncRNA that suppresses *BDNF* expression. It is well-established that BDNF is released from primary afferents [33] binding to TrkB in spinal cord neurons where it regulates synaptic plasticity important for chronic pain [11]. Our findings suggest that *BDNF*-*AS* could be an unappreciated suppressor of BDNF expression in females, a relevant finding because the role of BDNF signaling is more prominent in male than in female rodents [38; 48] and in male but not female human spinal cord [12].

At the level of transcription factor footprints, EGR1/3 and SP showed higher abundance in female accessible DNA in hDRG. In agreement with this, *EGR1* expression was higher in females. EGR1 is a transcription factor classified as immediate early gene. EGR1-driven transcriptional programs are activated by stimuli like cytokines, growth factors, oxidative stress, hypoxia and injury [85]. Further, Egr1 has an important role in neuronal plasticity in the adult nervous system [15]. A recent study found that the expression of *Egr1* is driven by female sex hormones and acts as a chromatin modifier in a sex-specific fashion [60], suggesting its pivotal role in controlling neuronal gene expression in females. Our data shows that EGR1 binding sites are found in genes like *JAK1* and *CX3CL1*, both of which are interferon associated genes known to induce pain through a peripheral mechanism in mouse models [2; 19]. This is also consistent with human thoracic vertebrectomy DRG data showing that interferon and JAK-STAT signaling are associated with neuropathic pain in female samples but not in males [59]. One inconsistency with the thoracic vertebrectomy data is that *EGR1* mRNA was more highly expressed in males with pain than females [59], but the impact of that difference likely depends on access to regulatory sites for the transcription factor. Also consistent with our findings, *SP4* has a higher expression in female tibial nerve compared to male [58]. SP4 regulates the transcription of pain-related genes [9] and, Sp4 knockdown reverses the hypersensitivity induced by persistent models of pain in mice, suggesting an important role of this transcription factor to pain [67]. Binding sites of other members of the SP (*SP1*/*SP5*) family were abundant in accessible chromatin in female hDRG, rendering these transcription factors potential therapeutic targets for treating pain in females since they are linked to neuropathies such as of chemotherapy-induced peripheral neuropathy [86].

In males, we observed a higher chromatin accessibility in *TRPV2/4* and Ca^2+^-related genes in hDRG, with the latter specifically in neurons. With respect to the former group of genes, many of these play important roles in pain but *TRPV4* is the only TRP channel that shows higher expression in pain samples from thoracic vertebrectomy pain patients in males versus females [59]. While we are not aware of any previous reports of sex differences in expression of Ca^2+^-related genes in hDRG, rodent studies demonstrate that some calcium-related genes including *Cacna1b* are upregulated in male DRG in a model of nerve injury, gene that showed higher accessibility in male hDRG in our data [71]. At the level of transcriptional programs, AP-1 transcription factors were predominant in male hDRG in both bulk and spatial ATAC-seq specifically in neurons. These data are in close agreement with increased JUN and FOS activated transcriptional cascades in DRGs from male patients with neuropathic pain in the thoracic vertebrectomy cohort [59]. We found that the promoters of genes such as *OSMR*, *NLRP3* or *CASP8* are predicted to be bound by these transcription factors. The abundance of binding sites for these genes in male open chromatin in the DRG increases the likelihood that they will be activated under conditions of inflammation or injury, which is consistent with the findings of the thoracic vertebrectomy cohort where OSM signaling was specifically upregulated in males [50; 59].

Collectively, our data demonstrate distinct baseline sex-specific chromatin accessibility patterns and transcriptional programs primed for activation by injury or inflammation in hDRG. Functional differences observed in female hDRG, such as increased accessibility of GABAergic, glutamatergic and interferon-related genes, could contribute to their heightened susceptibility to chronic pain. Conversely, higher accessibility of calcium signaling genes in males might underlie distinct pain mechanisms. These findings highlight the importance of considering sex as a biological variable in chronic pain research and potentially pave the way for the development of sex-specific therapeutic strategies.

## Limitations

One important limitation of our work is that we did not perform multi-ome analysis from the same cell, making it difficult to test for direct correlation of gene expression and chromatin accessibility. Further, our sample size is relatively small. This can be improved with additional studies, which should include samples from individuals with chronic pain. We note, however, that we could achieve our primary objective of detecting sex differences, many of which were replicated with two independent methods. Additional experiments like CUT&Tag sequencing [29] can be used to gain insight into the epigenetic landscape in the hDRG. Our work with organ donor DRGs shows that tissue quality is sufficient to enable these future studies.

## Supporting information

ATAC seq data files

## Acknowledgements

We acknowledge Yeunhee Kim, Manager of Genome Center at the University of Texas at Dallas for technical support with library prep for sequencing. We acknowledge Jennifer Garbarino, Andrés Hernández-Oliveras, Abhira Ravirala and Zawge Yohannes Daniel for the technical support to this project. The authors thank Anna Cervantes, Geoffrey Funk, and Peter Horton at the Southwest Transplant Alliance. The authors are grateful to the organ donors and their families for their gift. Data availability: Data in this paper will be uploaded to SPARC Portal. This work was supported by the National Institutes of Health grant U19NS130608 and R01NS111929 to Theodore J. Price.

