## supplementary figures for "Epigenomic landscape of the human dorsal root ganglion: sex differences and transcriptional regulation of nociceptive genes"

**Running title: epigenome of the human DRG**

Úrzula Franco-Enzástiga^1^, Nikhil N. Inturi^1^, Keerthana Natarajan^1^, Juliet M. Mwirigi^1^, Khadija Mazhar^1^, Johannes C.M. Schlachetzki^2^, Mark Schumacher^3^, Theodore J. Price^1,*^

^1^ Center for Advanced Pain Studies, School of Behavioral and Brain Sciences, University of Texas at Dallas, Richardson, Texas 75080.

^2^ Department of Cellular and Molecular Medicine, University of California, San Diego, 9500 Gilman Drive, La Jolla, CA 92093-0651, USA.

^3^ Department of Anesthesia and Perioperative Care and the UCSF Pain and Addiction Research Center, University of California, San Francisco, California, 94143 USA.

^*^Correspondence to:

Theodore J Price PhD

800 W Campbell Rd

BSB14.102

Richardson TX 75080

USA

+1 972-883-4311

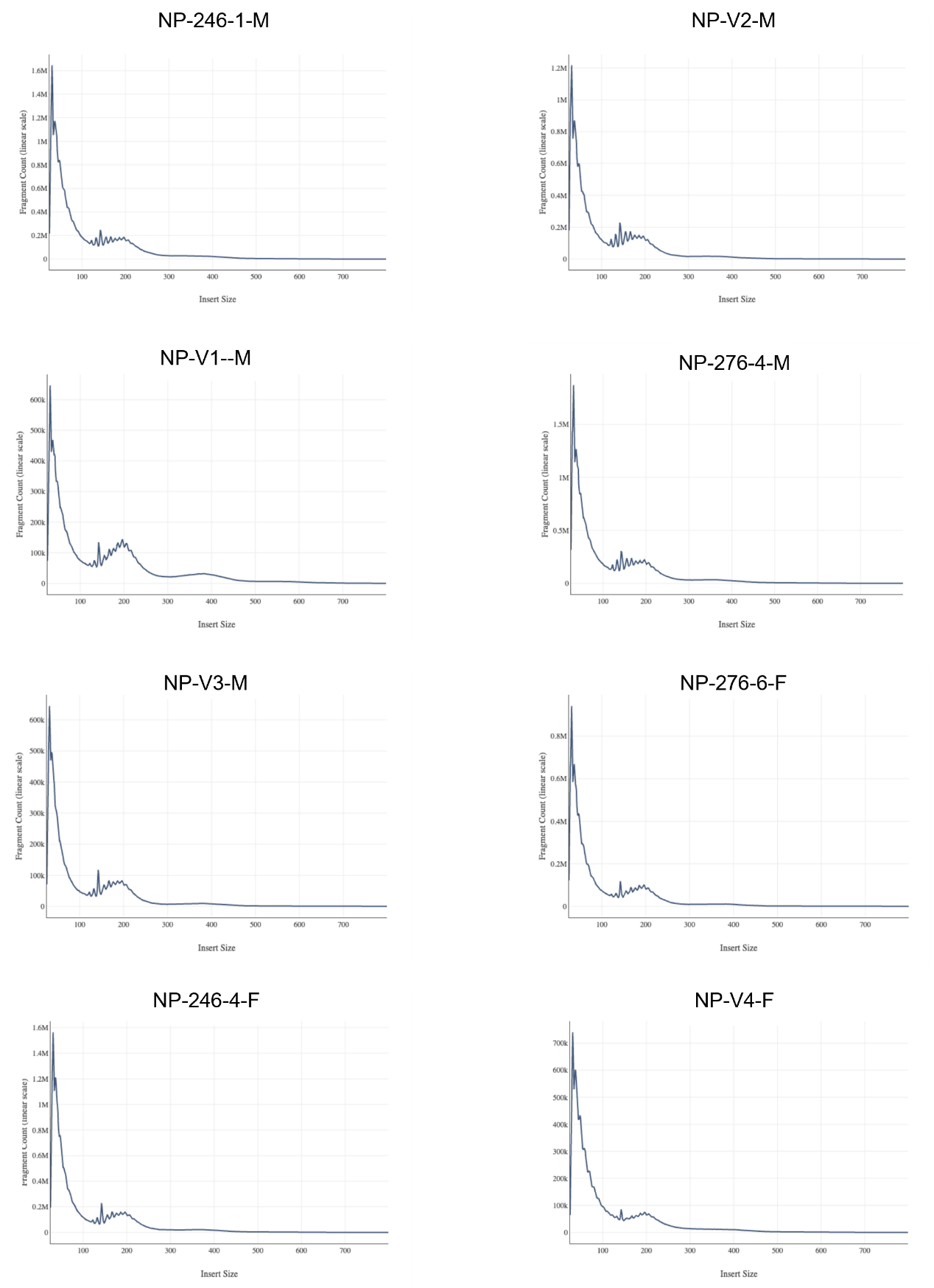

Fig. S1. Fragment size distribution plots for hDRG spatial ATAC-seq experiments. Nucleosome free regions are represented by the first peak in the fragment distribution plots. Peaks closer to 200 bp belong to mono-nucleosomes. Peaks around 400 bp represent di-nucleosomes.

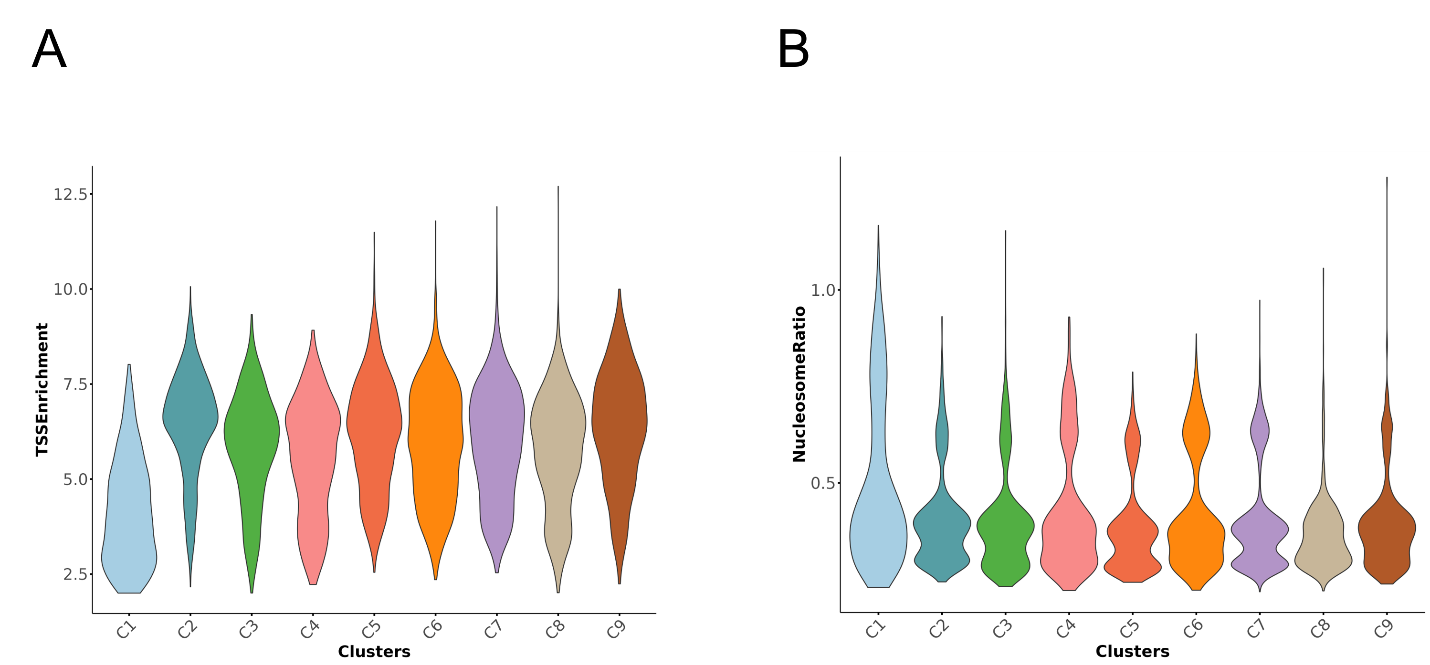

Fig. S2. A) TSS enrichment values and B) nucleosome ratio per barcode in spatial ATAC-seq clusters in hDRG samples.

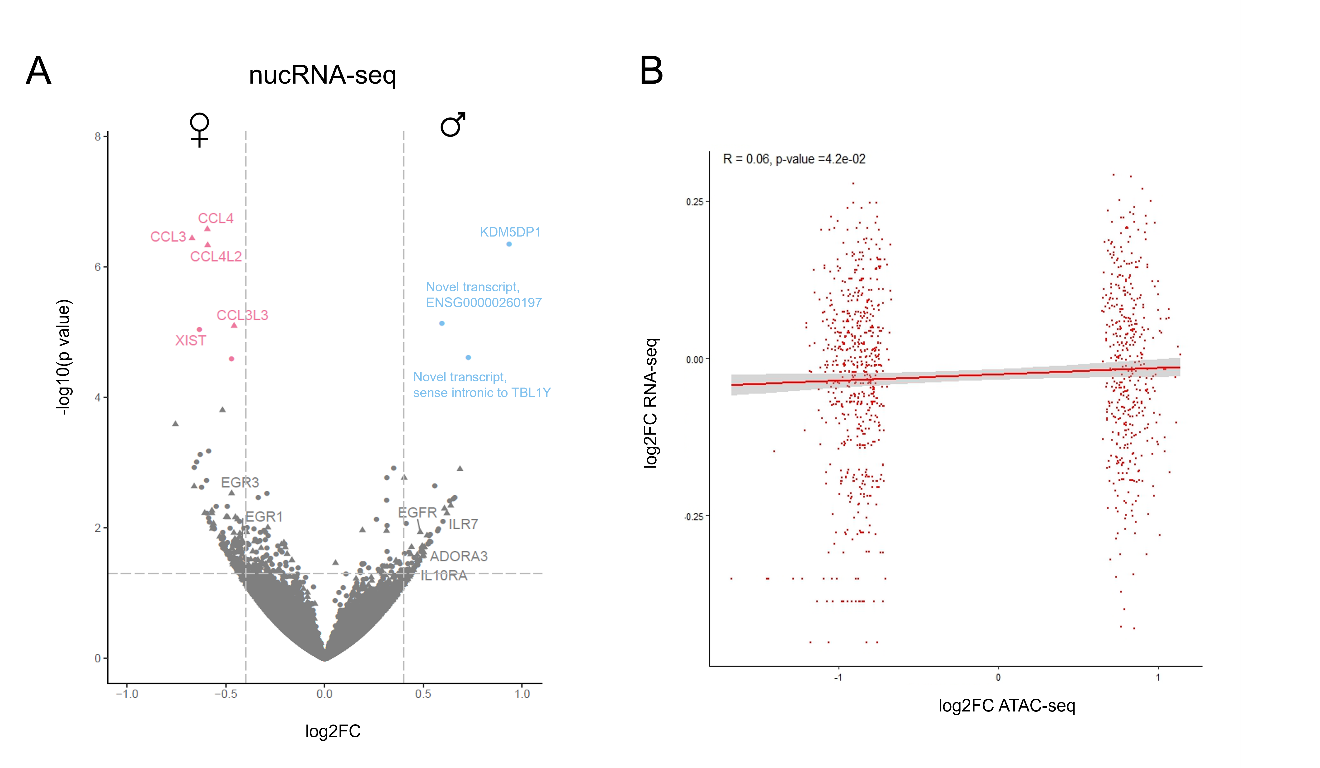

Fig. S3. Identification of differentially expressed genes (DEGs) in bulk nucRNA-seq from female and male hDRG. A) Volcano plot of all DEGs between sexes in hDRG (N = 7, 4 male (blue), and 3 females (pink) identified with DESeq2 (padj val <0.05; Fold change >1.33). Protein-coding genes are depicted with triangles and non-protein-coding genes with circles. B) The correlation plot (r) represents the Pearson correlation across DARs log2CF between female and male ATAC-seq signal (x axis) in gene bodies vs the log2FC RNA-seq gene expression between sexes (y axis).

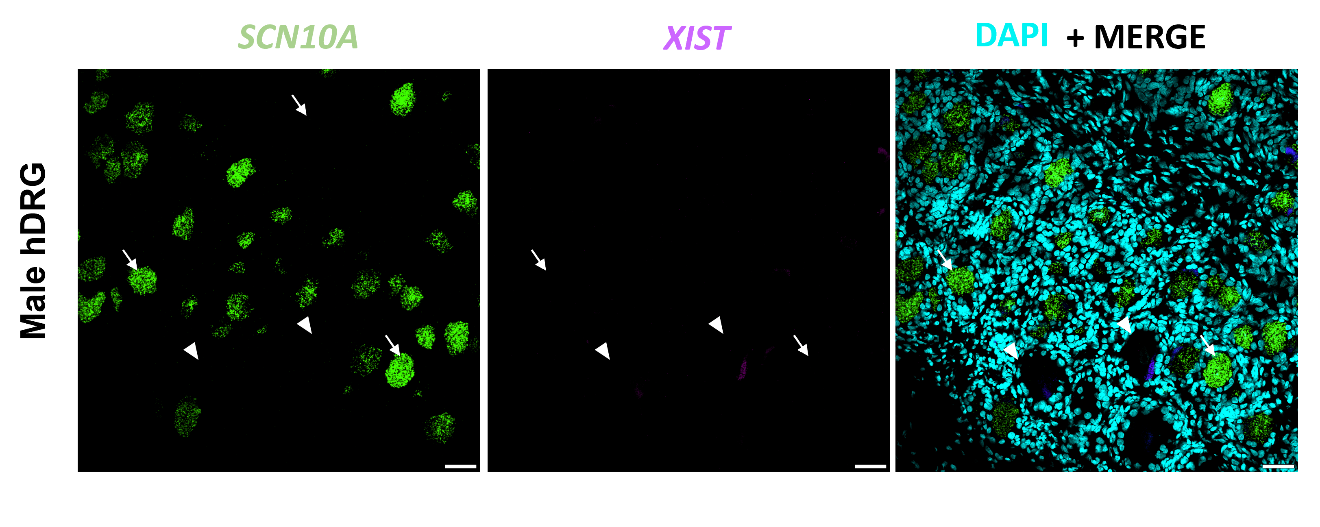

**Fig. S4**. *XIST* is barely detected in male hDRG. Representative *in situ* hybridization images of *XIST* (pink) in *SCN10A* (+) (green) and *SCN10A* (-) neurons in male hDRG (20X images). Scale bar – 50 µm. *SCN10A* positive neurons are indicated with an arrow. *SCN10A* negative neurons are indicated with an arrowhead.
